# Assessment of environmental and occupational exposure while working with multidrug resistant (MDR) fungus *Candida auris* in an animal facility

**DOI:** 10.1101/486100

**Keywords:** biocontainment, biosafety level 2, *Candida auris*, real-time PCR

## Abstract

In less than a decade since its identification in 2009, the emerging fungal pathogen *Candida auris* has become a major public health threat due to its multidrug resistant (MDR) phenotype, high transmissibility, and high mortality. Unlike any other *Candida* species, *C. auris* has acquired high levels of resistance to an already limited arsenal of antifungals. As an emerging pathogen, there are currently a limited number of documented murine models of *C. auris* infection. These animal models use inoculums as high as 10^7^-10^8^ cells per mouse, and the environmental and occupational exposure of working with these models has not been clearly defined. Using real-time quantitative PCR and culture, we monitored the animal holding room as well as the procedure room for up to six months while working with an intravenous model of *C. auris* infection. This study determined that shedding of the organism is dose-dependent, as detectable levels of *C. auris* were detected in the cage bedding when mice were infected with 10^7^ and 10^8^ cells, but not with doses of 10^5^ and 10^6^ cells. Autoclaving bedding in closed micro-isolator cages was found to be an effective way to minimize exposure to animal caretakers. We found that tissue necropsies of infected mice were also an important source of potential source exposure to *C. auris*. To mitigate these potential exposures, we implemented a rigorous “buddy system” workflow and a disinfection protocol that uses 10% bleach followed by 70% ethanol and can be used in any animal facility when using small animal models of *C. auris* infection.

## INTRODUCTION

Since its identification in 2009, the emerging fungal pathogen *Candida auris* has become a major public health threat, due to its status as a multidrug resistant (MDR) fungus with high levels of resistance to an already limited arsenal of antifungals, such as the azoles and polyenes.^(1–4)^ Minimal inhibitory concentrations (MICs) for antifungals used against *C. auris* have been reported to be as high as 256 mg/L for fluconazole and > 2 mg/L for amphotericin B.^(1–5)^ The Centers for Disease Control (CDC) has documented that *C. auris* behaves more like transmissible bacterial multidrug resistant organisms (MDROs) such as methicillin resistant *Staphylococcus aureus* (MRSA) than any other *Candida* species.^(6)^ Of particular concern is that the unpredictable antifungal resistance profile of *C. auris* is also showing evidence of reduced susceptibility to the echinocandins, a third class of antifungals.^(4)^ *C. auris* causes invasive bloodstream infections with reported mortality rates as high as 30-60% in Venezuela and India, and most recently, a reported mortality rate of up to 78% in Panama.^(7–9)^ Although the origin of *C. auris* is unknown, infections with this organism have been documented on five continents in at least 18 countries, including the United States.^(7, 8, 10–18)^

*C. auris* has a predilection for the skin, making transmission and colonization from person to person quite easy ^(6)^. *C. auris* has been found to be alarmingly persistent in the environment, and has been reported in hospital and nursing home settings. ^(19, 20)^ A recent study at the Louis Stokes Veterans Affairs Medical Center in Cleveland, Ohio found that *C. auris* exhibits a greater propensity to persist for up to 7 days on surfaces than any other *Candida* species. Surfaces that tested positive included moist surfaces, such as hospital sinks and shower drains, and dry surfaces like bed rails, tables, and call buttons. ^(21)^ *C. auris* is often misidentified as *Candida haemulonii*, and without specialized laboratory methodologies such as Matrix-Assisted Laser Desorption Ionization Time of Flight Spectrometry (MALDI-TOF), diagnosis is delayed, leading to mortality rates as high as 35%.^(16, 19, 22)^

The current animal models of disseminated *C. auris* infection use concentrations that range from 10^6^-10^8^, along with the immunosuppressive agent cyclophosphamide. These animal models are being used to test new alternative therapeutics, including a small molecule antifungal agent called APX001A derived from the pro-drug APX001 that inhibits the fungal protein Gwt1 (glycosylphosphatidylinositol-anchored wall transfer protein 1). A novel echinocandin called rezafungin is also being tested in these animal models as a potential therapeutic to combat *C. auris*.^(23–26)^ Because of its multidrug resistance, environmental persistence, and the large inoculum used when working with animal models infected with *C. auris*, the risk of an occupational exposure during experiments for researchers and animal care staff is still unclear.

In this study, immune-competent and neutrophil-depleted mice were infected with the *C. auris* strain M5658 that belongs to the South Asia clade. The concentrations of *C. auris* used to infect mice ranged from 10^5^-10^8^ inoculum per animal to determine potential for exposure.^(27)^ The animal holding room and procedure room that we worked in were monitored to determine whether there was a risk for exposure during cage changes, animal infections, and animal procedures such as necropsies. As part of this study, a rigorous “buddy system” workflow, disinfection protocol and safety procedures were developed to prevent accidental exposures.

## METHODS

### Animal Use Ethics Statement

Animal experiments described in this study were approved by the Wadsworth Center’s Institutional Animal Care and Use Committee (IACUC) under protocol #18-450. The Wadsworth Center complies with the Public Health Service Policy on Humane Care and Use of Laboratory Animals and was issued assurance number A3183-01. The Wadsworth Center is fully accredited by the Association for Assessment and Accreditation of Laboratory Animal Care (AAALAC).

### The “Buddy System” Workflow: Handling and Cleaning of Infected Cages in Animal Holding Room

*C. auris* is recommended by the Centers for Disease Control and Prevention (CDC) to be handled at Biosafety Level 2 (ABSL-2). However, due to the limited knowledge of its environmental exposure risk in an animal research setting, the organism was handled at Animal Biosafety Level 2 (ABSL-2) with Animal Biosafety Level 3 (ABSL-3) containment practices that included enhanced Personal Protective Equipment (PPE) and a strict no open handling policy. Staff entering the animal facility used standard PPE that included booties, gloves, bouffant, face mask, and a gown. Before entering the animal holding room, additional PPE was donned which included a second pair of booties, gloves and a snap-front gown (Valumax, New York, NY).

Mice infected with *C. auris* were housed in locked micro-isolator cages (Tecniplast, West Chester, PA) with a negative air flow set to 51%, RH% 47, 75 ACH. Micro-isolator cages also provided a way to autoclave bedding and water bottles as a single unit after use, thus minimizing exposure to animal caretakers. A “buddy system” was implemented such that one person could hand the clean caging and other supplies to a second person performing the cage changes inside of the biosafety cabinet (Figure 1). This allowed for a “clean” to “dirty” workflow with minimal disruption of the laminar airflow of the biosafety cabinet. When there was a need to remove hands from the biosafety cabinet, the exterior gloves were doffed inside the cabinet and a new pair was donned. Blue pads placed inside of a biosafety cabinet (Kendall Healthcare Products, Mansfield, MA) were sprayed with a 10% bleach solution. It is important to note that the 10% bleach solution was made fresh daily as recommended by the CDC, since quaternary ammonia products routinely used for disinfection in animal research settings are not effective against *C. auris*.^(4, 28)^

**Figure 1.**
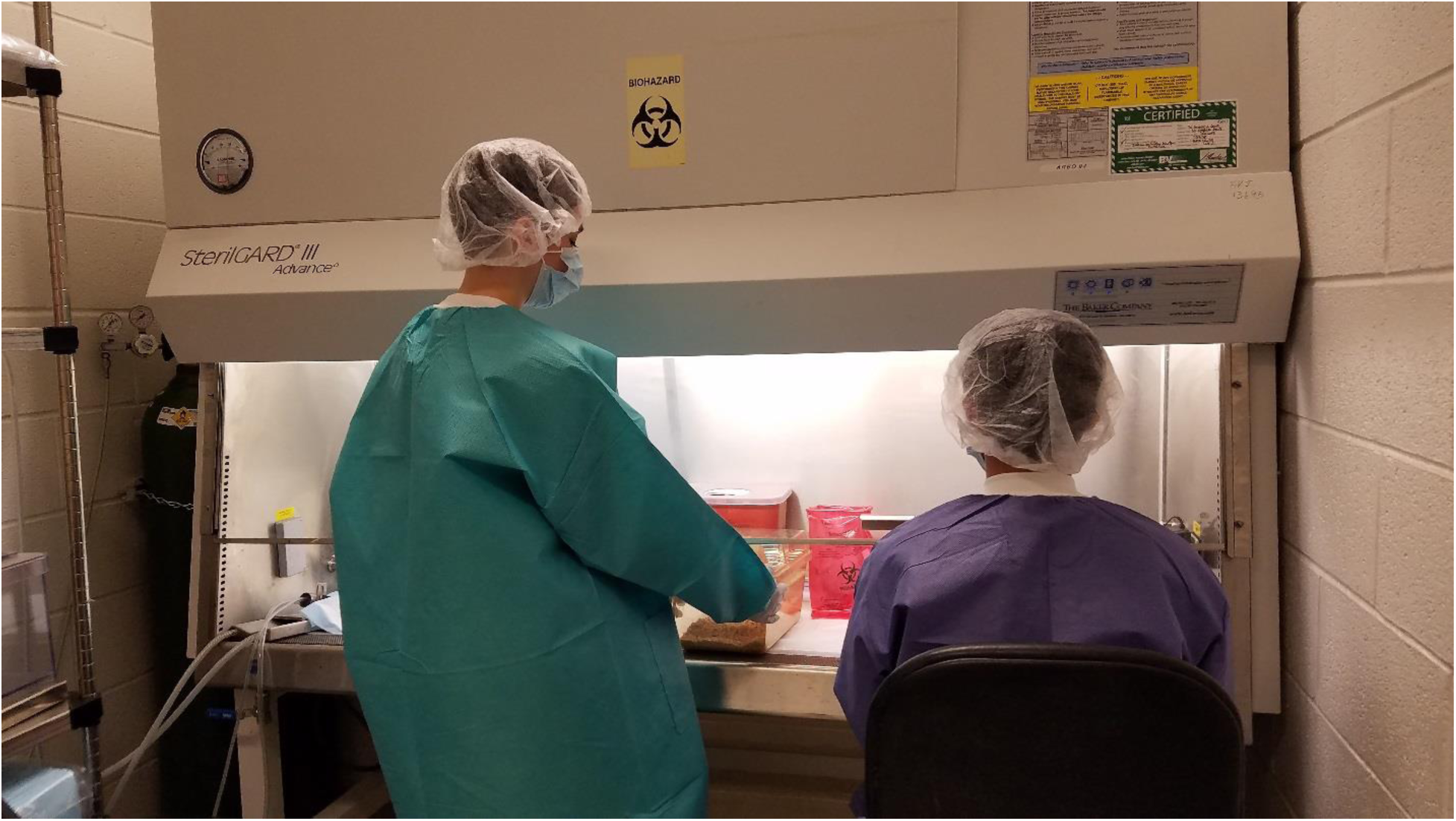
The “buddy system” work flow. A “buddy system” was implemented, such that one person hands over clean caging and other supplies to a second person performing the procedures inside of the biosafety cabinet. This allows for a “clean” to “dirty” workflow with minimal disruption of the laminar airflow of the biosafety cabinet.

To change infected animals from cages, the in-use micro-isolator cage and an empty clean cage were placed inside of the biosafety cabinet on top of the blue pads. Mice were carefully transferred to the new cage. Cage bedding that fell into the biosafety cabinet was collected by the blue pads sprayed with 10% bleach. Food was added to the new cages from a stock strictly kept inside of the biosafety cabinet. Before removal from the biosafety cabinet, cages were wiped down with 10% bleach by the individual inside the cabinet, followed by a wipe down of 70% ethanol by the individual outside the cabinet. At the end of cage changes, the blue pads were rolled up and placed into a biohazard waste bag. The biosafety cabinet and contents were sprayed down with a 10% bleach solution and allowed to sit for 5 minutes, followed by wipe down with 70% ethanol (Figure 2). Before leaving the room, the exterior booties, gloves and gown were doffed. Dirty micro-isolator cages were sent to be autoclaved as a single sealed unit with bedding and water bottle still inside to reduce potential occupational exposures to animal caretakers during the transport of cages to the autoclave.

**Figure 2.**
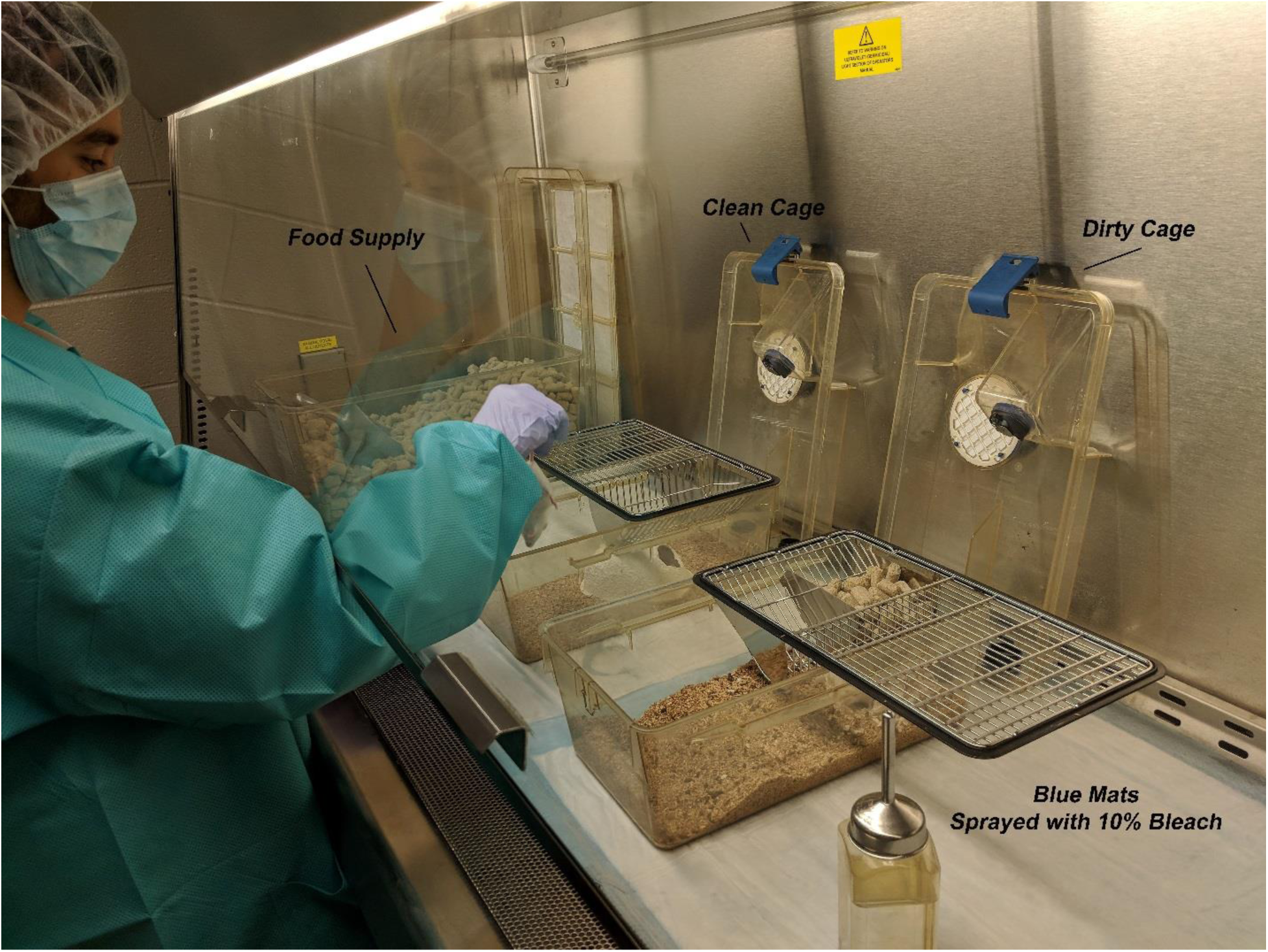
Workflow for cage changes during *C. auris* experiments. The in-use micro-isolator cage and an empty clean cage were placed inside of the biosafety cabinet atop the blue pads that were sprayed with 10% bleach. Mice were carefully transferred to the new cage and food was added to the new cages. The food stock was kept strictly kept inside of the biosafety cabinet. Before removal from the biosafety cabinet, cages were wiped down with 10% bleach by the individual inside of the cabinet, followed by 70% ethanol by the individual outside of the cabinet. At the end of cage changes, the blue pads were rolled up and placed into a biohazard waste bag. The biosafety cabinet and contents were sprayed down with a 10% bleach solution and allowed to sit for 5 min and wiped. Additionally, the cabinet was cleaned with a 70% ethanol solution.

### Transferring Cages from Holding to Procedure Room

Cages with infected animals were transferred onto a laboratory cart that was placed outside of the holding room. The cart contained dry blue pads that were not sprayed with 10% bleach; the cages were placed on each shelf of the cart. Before leaving the holding room, exterior gloves, booties and gowns were removed and placed in a biohazard waste bin. The cages were then transferred from the holding room to the procedure room. When entering the procedure room, clean outer booties, outer gloves, and snap-front gown were donned. The cart with the isolator cages as well as any supplies was kept on the “clean” side of the room, indicated by a line on the floor.

### Animal Handling during Weight Measurements

Dry blue pads were placed inside of a biosafety cabinet to cover the surfaces and cages were placed on top. To monitor daily weights throughout the experimental trial, mice were placed into a sterilized container that contained airholes and weighed using a laboratory balance. After the last mouse in each cage was weighed, the cage was sealed and wiped down with 10% bleach by the person inside of the cabinet, followed by a wipe with 70% ethanol by the person outside of the cabinet, before returning cages to the cart. This was repeated until all mice had been weighed. At the end of workflow, if no other procedures were done, the blue pads in the biosafety cabinet were rolled up and placed in a biohazard waste bag. The biosafety cabinet, scale, weigh chambers, and all contents were sprayed with a 10% bleach solution. After 5 minutes, the surfaces were wiped and then cleaned with a 70% ethanol solution (Figure 3).

**Figure 3.**
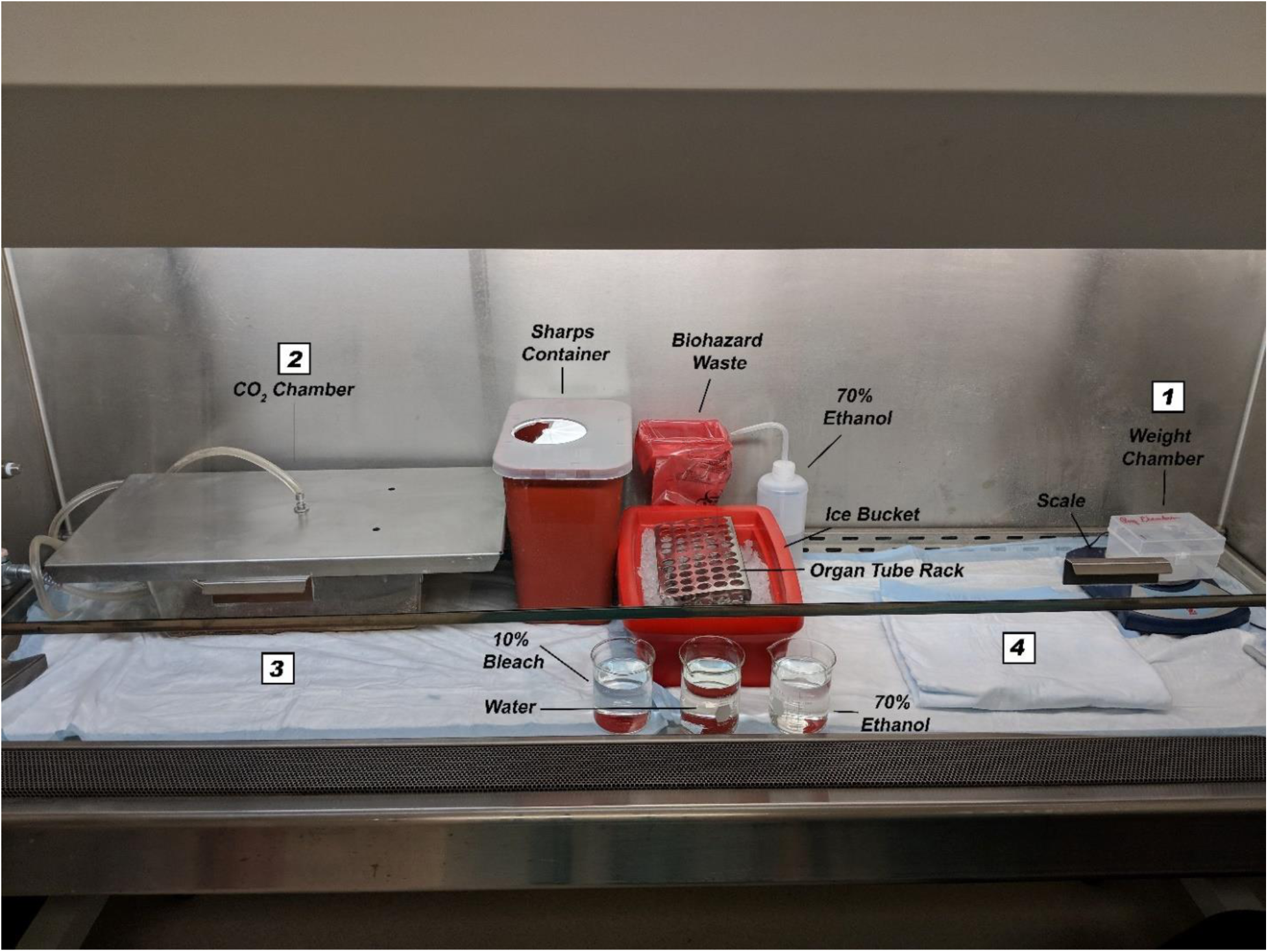
Biosafety cabinet setup within the procedure room. (1) On the right-hand side of the biosafety cabinet, mice were weighed within the chamber on the scale. Daily weights were taken for all mice in the study before moving on to the next step. (2) Mice were euthanized in the CO_2_ chamber on the left-hand side, as the CO_2_ source was only available on that side of the biosafety cabinet. (3) Cardiac punctures were performed directly after CO_2_ euthanasia. Procedure needles were disposed of in the sharps container. Cervical dislocation was done as a secondary method of animal euthanasia. (4) The mice were transferred to the folded blue mat on the right. 70% ethanol was sprayed on the mouse’s abdomen prior to making a midline incision for organ collection. Organs were collected and placed inside labeled tubes containing 2mL HBSS on the organ tube rack. When necropsy was finished, the carcass was placed into a sealable biohazard bag. Tools were rinsed in the following order: 10% bleach, water, then 70% ethanol between each mouse. The folded blue mat was switched to a clean side between mice. Any waste generated throughout necropsies was put into the biohazard bag in the back of the biosafety cabinet. At the end of workflow, the blue pads were rolled up and placed in the biohazard waste bag. The cabinet and all contents were sprayed with 10% bleach and allowed to sit for five min. After five min all contents and the cabinet were wiped and then cleaned using 70% ethanol.

### Animal Infections and Necropsies

Prior to *C. auris* infection, 100 µg of the monoclonal antibody RB6-5C8 that depletes Ly6G neutrophils (Thermo Fisher, San Diego, CA) was administered by intraperitoneal (i.p.) injection using a 25G needle (BD, Franklin Lakes, NJ). Twenty-four hours later, 100 µl of *C. auris* was administered via tail vein (i.v.) injection at concentrations ranging from 10^5^-10^8^ cells. Prior to infection, mice were initially anesthetized in an isoflurane chamber located outside of the biosafety cabinet. Once the mouse was anesthetized, the individual outside the biosafety cabinet placed it inside the cabinet. The individual with the hands inside the cabinet then placed the mouse into an illuminated mouse restrainer (Braintree Scientific, Braintree, MA) and placed a nose cone on the mouse. After the injection, the mice were transferred back into the cage located inside of the biosafety cabinet. The injection syringe was immediately placed in a sharps container to prevent needlestick accidents. When all mice in a particular group had been injected, the cage was wiped with 10% bleach by the individual inside of the cabinet, followed by a wipe with 70% ethanol by the individual outside of the cabinet. This procedure was repeated until all groups had been injected. To collect organs by necropsy, mice were euthanized in a CO_2_ chamber located inside the biosafety cabinet, and death was confirmed by cervical dislocation. Blood was collected via cardiac puncture using a 26G needle, and spleen, liver, kidney, stomach, heart, brain and lungs were collected and placed in labeled 5 ml tubes (CELLTREAT, Pepperell, MA) containing 2 ml of HBSS with no dye on ice. Organs were ground through a 70 µm cell strainer (CELLTREAT, Pepperell, MA) using a pestle (CELLTREAT, Pepperell, MA), and twenty microliters (20 µl) of the resulting homogenate were plated on Sabouraud Dextrose Agar plates with antibiotics in triplicate and incubated at 37°C for 24-hr for colony forming units (CFU). When there was a need to remove hands from inside the biosafety cabinet, the exterior gloves were doffed and a new pair donned before continuing to work. Glove changes were also done in between handling of different treatment groups to avoid cross-contamination. Surgical tools were sanitized with 10% bleach, 70% ethanol, and water between mice. The carcasses were placed into a sealable biohazard bag. At the end of the procedure, blue pads were rolled up and placed into a biohazard bag. All contents of the biosafety cabinet including the ice bucket, tubes where organs were collected, and surgical tools were sprayed with 10% bleach, allowed to sit for 5 min, and then wiped down with 70% ethanol. The biohazard bag containing the mouse carcasses was placed into a second sealable biohazard bag and then frozen prior to final disposal by incineration. The biohazard bags containing blue pads and any waste were placed into a biohazard waste bin. The biosafety cabinet was wiped down with 10% bleach and then cleaned with 70% ethanol.

### Return of Animals to the Holding Room and Tissue Transport for Processing

When mice were returned to the holding room, the blue pads from the cart were disposed of in a biohazard bag and the cart wiped with 10% bleach followed by 70% ethanol. The collected and ground organ samples were moved from the ice bucket and placed inside a specimen transport box (Nalgene, Rochester NY). The transport box was also sprayed with 10% bleach for 5 min followed by 70% ethanol, and then placed on the clean side of the room to await transfer to the laboratory.

### Environmental Monitoring of *C. auris* in Holding and Procedure Rooms by Surface Swab testing and real-time quantitative PCR (qPCR)

Prior to doing any work with *C. auris,* baseline surface swabs in both the holding and procedure rooms were tested. We also performed surface swab testing after any procedure such as cage changes or necropsies and after disinfection with 10% bleach followed by 70% ethanol. Swabs were taken on selected surfaces; for example, in the animal holding rooms where routine cage changes were done, we tested the air flow nozzle of the micro-isolator cages, the floor near the biosafety cabinet, the interior door handle, the exterior door handle, and the cart used to transport mice between the holding and procedure rooms. The floor of the room, door knobs, and the transport cart were also monitored to identify potential cross-contamination that might have arisen from improper PPE use or clean-up procedure. Surfaces were swabbed using a sponge-stick (3M Health Care, St. Paul, MN), the sponges placed back into their respective bags, and the handles removed. The bag was folded four to five times and sealed, taking care that only the sponge touched the inside of the bag during sampling. Surfaces were again wiped with 10% bleach followed by 70% ethanol to remove residue left by the sponge-stick. In our laboratory, the sponges were processed immediately, but could be stored at 4°C in the sealed bag for up to a week. The protocol for sponge processing was done as previously described by Leach et al., who developed the method for rapid identification of *C. auris* by PCR.^(27)^ Briefly, in a Whirl-Pak bag (Nasco, Fort Atkinson, WI), 45 ml of 1x PBS with 0.02% of Tween 80 was added to the swabbing sponge and placed in a stomacher 400 circulator (Laboratory Supply Network, Inc., Atkinson, NH) for 1 min at 260 rpm. The liquid was centrifuged at 4,000 rpm for five min and the supernatant was decanted, leaving approximately 2 ml of liquid in the bottom of the tube. One milliliter of sample was used for plating and the other 1ml was used for DNA extraction. The sample was centrifuged at 13,000 rpm for five min and the collected pellet was washed twice, then resuspended in 48 µl of PBS-BSA. The 48 µl was then transferred to a sterile screw cap tube (Heathrow Scientific, Vernon Hills, IL) containing two 3 mm glass beads and 2 μl of bicoid inhibition control plasmid DNA (5 × 10^−3^ ng/μl).^(27)^ The tubes were then homogenized using a Vortex Genie in 10 min intervals followed by a 2 min incubation on ice for a total of 50 min of homogenization.

After collection, swabs were analyzed via real-time quantitative (qPCR) designed specifically for the detection of *C. auris* DNA.^(27)^ Samples were analyzed by qPCR immediately, or stored at −20°C. The qPCR targets the internal transcribed spacer 2 (*ITS*2) gene of *C. auris*.^(27)^ Briefly, as previously described by Leach et al., who developed the real-time PCR assay, the PrimerQuest program (Integrated DNA Technologies, Coralville, IA) was used to design primers and a probe from this region. Primers and a probe were also designed for a *bicoid* gene serving as an inhibition control. The sequences for the primers and probes for *C. auris ITS2* gene were as follows: V2424F (CAURF), 5′-CAGACGTGAATCATCGAATCT-3′; V2425P (CAURP), 5′-/56-carboxyfluorescein (FAM)/AATCTTCGC/ZEN/GGTGGCGTTGCATTCA/3IABkFQ/-3′; and V2426 (CAURR), 5′-TTTCGTGCAAGCTGTAATTT-3′. For the *bicoid* gene, primers and probes were as follows: V2375 (BICF), 5′-CAGCTTGCAGACTCTTAG-3′; V2384 (BICP), 5′/Cy3/AACGCTTTGACTCCGTCACCCA/3IAbRQSp/-3′; and V2376 (BICR), 5′-GAATGACTCGCTGTAGTG-3′. All primers and probes were obtained from Integrated DNA Technologies.

Each infected tissue sample was tested in duplicate in 20-μl volumes using an optical 96-well reaction plate. Each reaction mixture contained 1× PerfeCTa Multiplex qPCR ToughMix (Quanta Biosciences), a 500 nM concentration of each *C. auris* primer (V2424 and V2426), a 100 nM concentration of each *bicoid* primer (V2375 and V2376), a 100 nM concentration each of *C. auris* (V2425P) and *bicoid* probe (V2384P), and 5 μl of DNA extracted either from *C. auris* or from the surveillance samples. Each PCR run also included 5 μl of positive extraction (*C. auris* M5658; 10^3^ CFU/50 μl) and positive amplification (*C. auris* M5658; 0.02 pg/μl) controls, as well as 5 μl of negative extraction control that contained reagents only and negative amplification control that contained sterilized nuclease-free water. To prevent any cross-contamination, a unidirectional workflow was followed that kept reagent preparation, specimen preparation, and amplification/detection areas separate. qPCR was performed in a room specifically designated for the amplification of *C. auris* DNA. The cycling conditions used on the ABI 7500 FAST were 95°C for 20 s, followed by 45 cycles of 95°C for 3 s and 60°C for 30 s. Based on receiver operating characteristic (ROC) curve analysis, a cycle threshold (*C*_*T*_) value of ≤37 was reported as positive and >37 was reported as negative for swabs. Similarly, a *C*_*T*_ value of ≤38 was reported as positive and >38 was reported as negative for sponges. Specimens were reported as inconclusive if PCR inhibition was observed for either swabs or sponges.

### Environmental Monitoring of *C. auris* in Holding and Procedure Rooms by Culture

While changing animals from their cages in the animal holding room or when performing necropsies in the procedure room, we placed opened Sabouraud Dextrose Agar plates with antibiotics (40 μg/ml streptomycin; 20 U/ml penicillin; 25 μg/ml chloramphenicol; 40 μg/ml gentamicin) inside the biosafety cabinet, as well as on the floor directly outside of the cabinet. At the completion of the work flow, these plates were incubated at 37°C for up to two weeks and checked daily to monitor growth of *C. auris.* These plates were used as a secondary measure to determine whether *C. auris* was aerosolized within the biosafety cabinet or whether any debris during animal handling such as cage bedding or tissue had a potential of contaminating the floor near the biosafety cabinet. Assessment of floor contamination was important, as the animal caretakers cleaned our floors on a weekly basis and required guidance as to whether to clean the floors with 10% bleach.

### Assessment of *C. auris* Shedding in Mice by Culturing of Oral and Anal Swabs

Oral and anal swabs were taken from neutrophil-depleted mice infected with *C. auris* and immune-competent control mice using Q-tips (Puritan, Guilford, ME). The Q-tips were placed in labeled 15 mL tubes, and each was submerged in 1 mL 0.02% PBS-Tween-80 and vortexed for 10 sec. The Q-tip was pressed against the sides of the tube as it was lifted out to remove as much moisture as possible. The Q-tip was discarded in a biohazard bag. The liquid collected was transferred to a labeled 2 ml screwcap tube. These samples were stored at 4°C for no longer than one week before processing was continued. The collected liquid was cultured in three ways: (1) 50 µl of the liquid was plated on Sabouraud Dextrose Agar plates with antibiotics, (2) 50 µl of the liquid was plated on Sabouraud Dulcitol Agar plates with 10% NaCl and antibiotics, and (3) 200 µl was inoculated in a Sabouraud Dulcitol Agar, 10% NaCl and antibiotics broth. Plates were incubated at 40°C for two weeks. Broth cultures were incubated in a shaker at 40°C for two weeks. Cultures that were positive were verified by PCR as previously described by Leach et al.(27)

## RESULTS

### Monitoring Animal Holding and Procedure Rooms that Contain Mice Infected with *C. auris*

Due to the persistence of *C. auris* in hospitals and nursing home settings, coupled with its resistance to antifungals, it was important to identify potential sources of exposure while working with intravenously (i.v.) infected mice. Mice were infected with *C. auris* at a range of 10^5^-10^8^ cells. At concentrations of 10^8^, *C. auris* was fulminant to neutrophil-depleted mice, as these animals succumbed to infection in less than 24 hr. At the lowest inoculum of 10^5^ cells, *C. auris* targeted multiple organs such as the spleen, liver, kidney, stomach, heart, lungs, and brain. Immune-competent mice exhibited 3.65×10^3^ cells per organ for spleen, 7.85×10^1^ cells per organ for liver, 2.80 ×10^2^ cells per organ for kidney, 1.07×10^3^ cells per organ for heart, and 4.30×10^1^ cells per organ for brain. No *C. auris* was detectable in the lungs and stomach. Neutrophil-depleted mice exhibited 1.81×10^4^ cells per organ for spleen, 1.12×10^2^ cells per organ for liver, 5.78×10^2^ cells per organ for kidney, 4.22×10^1^ cells per organ for stomach, 1.56 ×10^3^ cells per organ for heart, and 1.70×10^3^ cells per organ for lungs. No *C. auris* was detected in the brain in these mice (Figure 4). During our infection experiments, we noted that in both neutrophil depleted and animals those with an intact immune system, the spleen and heart are the organs most targeted by *C. auris* when introduced by tail vein injection.

**Figure 4.**
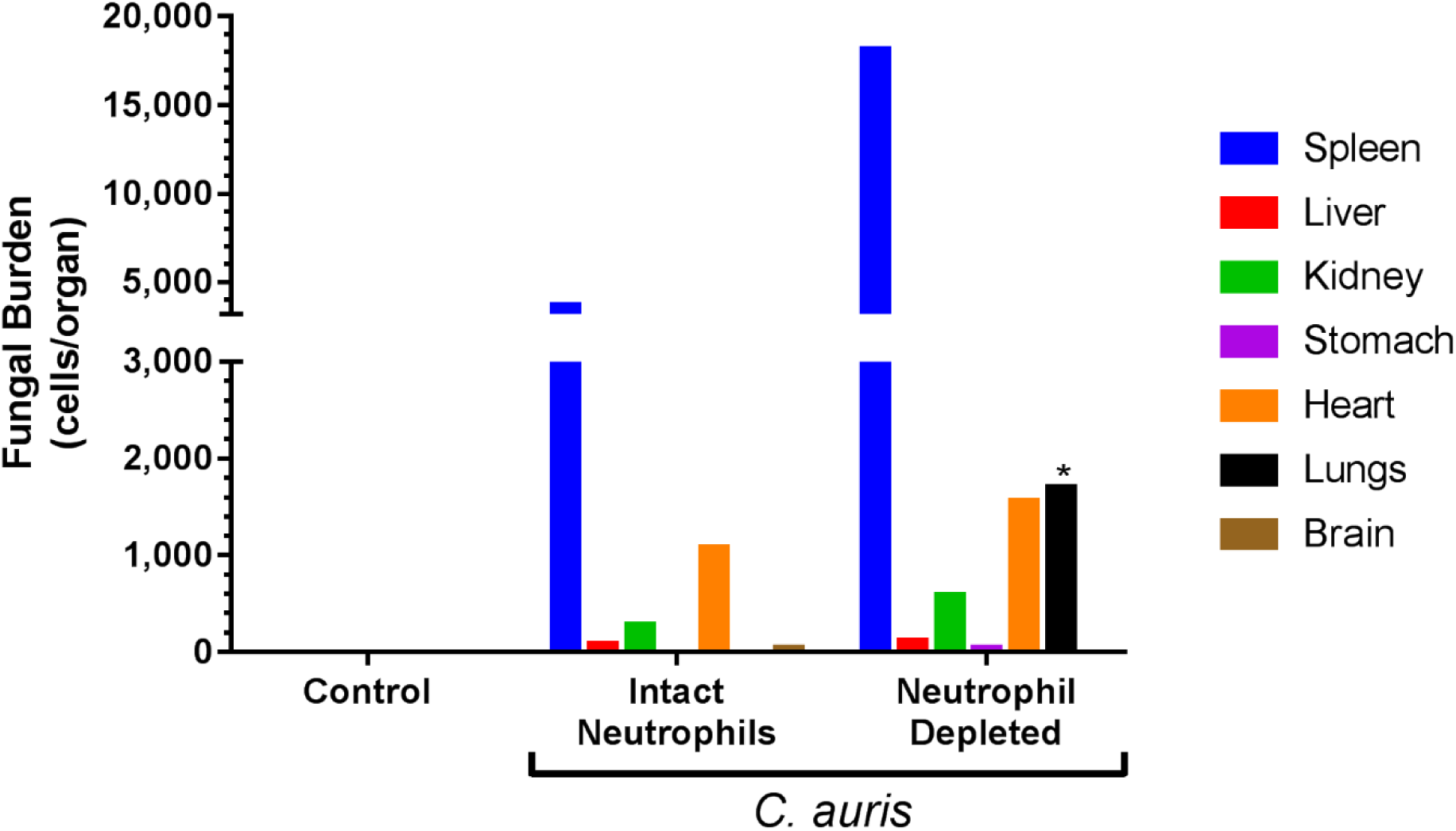
Organ fungal burdens in *C. auris* mouse model of infection. Organ fungal burden in mice infected with 10^5^ *C. auris* by tail vein injection. Neutrophil-depleted and control mice were assessed five days post infection. Neutrophil-depleted mice were found to have additional fungal burden in the lungs (black bar). Statistical significance in the lung p <0.05. Data represent mean cells per organ; n = 4 mice per group.

To assess the potential sources of *C. auris* contamination and exposure to researchers or animal caretakers during the study, both the animal holding and procedure rooms were continuously monitored by surface swab testing, qPCR and culture. The room surfaces monitored for *C. auris* were chosen based on their direct or indirect contact with infected mice and the potential for exposure to lab personnel. We swabbed the air flow nozzle of the micro-isolator cages, the floor near the biosafety cabinet, the interior door handle, the exterior door handle, and the cart used to transport mice between the holding and procedure rooms. The air flow nozzle of the mouse rack was particularly important as it served as an indicator of whether *C. auris* could be horizontally transmitted in the animal room, as has been described for *C. albicans*.^(29)^ Table 1 shows 5 days of surface swab data that were done in replicates to ensure reproducibility. To eliminate false negative results, during the qPCR a bicoid plasmid was added to the samples as positive internal control. The results reveal that at all concentrations of infection (10^5^-10^8^) of *C. auris* that we worked with, there was no horizontal transmission. Our buddy system and disinfection practices were effective, as there were no detectable levels of *C. auris* in the monitored animal holding room, including the floor of the room, door knobs, or the transport cart. We also monitored whether bedding that might potentially fall on the floor during cage transfers was a potential source of exposure by placing opened Sabouraud Dextrose Agar plates with antibiotics the floor directly outside of the cabinet. The results showed no growth after two weeks (data not shown).

**Table 1.** Monitoring of Animal Holding Room by qPCR

|  | Target | Baseline | Day 1 | Day 2 | Day 3 | Day 4 | Day 5 |
| --- | --- | --- | --- | --- | --- | --- | --- |
| **Mouse Rack Nozzle** | *C. auris* | (-) | ND | (-) | (-) | (-) | (-) |
|  | Bicoid Plasmid | 26.87±0.07 | ND | 28.16±0.18 | 28.46±0.52 | 26.20±1.86 | 26.51±1.05 |
| **Floor near BSC** | *C. auris* | (-) | ND | (-) | (-) | (-) | (-) |
|  | Bicoid Plasmid | 28.03±0.39 | ND | 28.07±0.14 | 29.83±0.28 | 26.04±0.67 | 27.09±0.17 |
| **Interior Door handle** | *C. auris* | (-) | (-) | ND | (-) | (-) | (-) |
|  | Bicoid Plasmid | 29.38±1.15 | 28.30±1.17 | ND | 29.54±0.83 | 29.07±0.34 | 27.74±0.30 |
| **Exterior Door handle** | *C. auris* | (-) | (-) | ND | (-) | (-) | (-) |
|  | Bicoid Plasmid | 28.63±0.61 | 27.57±1.26 | ND | 27.88±0.08 | 29.23±0.10 | 27.88±0.85 |
| **Lab Cart 3-tier** | *C. auris* | (-) | (-) | (-) | (-) | (-) | (-) |
|  | Bicoid Plasmid | 27.07±0.90 | 27.58±0.06 | 28.93±.25 | 28.37±0.04 | 29.26±0.61 | 29.01±0.20 |

Monitoring of the animal facility holding room where daily cage changes are done. Ct values show (-) negative detection of *C. auris* DNA by qPCR. Bicoid positive internal control shows positive Ct values. N.D. denotes not determined.

Monitoring of the procedure room was also assessed. This room was particularly important, as blood collection, centrifugation and necropsies were performed there. Surfaces such as the isoflurane nose cone, floor near the biosafety cabinets, chair backs, interior and exterior of the sink faucets and spigots, and interior and exterior door handles were tested. The results, summarized in Table 2, reveal that there was no *C. auris* in the animal holding room itself. Several replicates show undetectable levels on the door handles, faucet spigot, sink, chairs and stockpots. Even items inside the biosafety cabinet, such as the isofluorane nose cone and sharps container, were negative. While the water bottle tip tested negative, samples of the cage bedding that held infected mice, the interior working surface of the biosafety cabinet where necropsies were performed, and exterior gloves all tested positive at certain *C. auris* concentrations (Table 3; Figure 5). The results shown in Table 3 reveal that at 10^7^, mouse bedding and exterior gloves had positive Ct values of 29.52 and 28.36, respectively. At the 10^8^ inoculum, mouse bedding, exterior gloves, and the working surface of the biosafety cabinet had positive Ct values of 28.90, 29.01, and 22.93, respectively, suggesting that at these concentrations, mice begin to shed *C. auris* into the bedding. No shedding into the bedding was found at inoculums of 10^5^ and 10^6^, as the PCR was negative.

**Figure 5.**
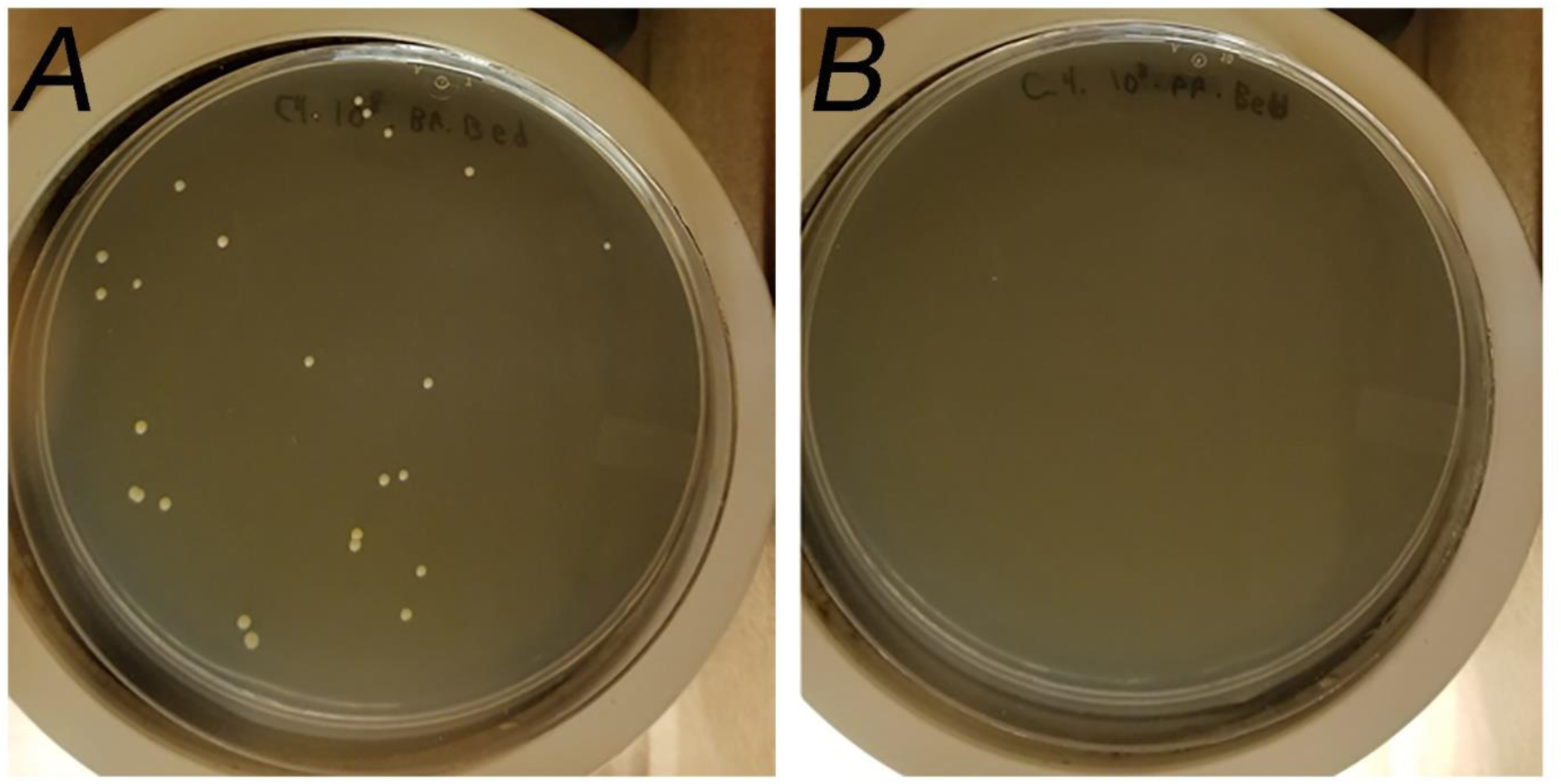
Autoclaving is an important step in disinfecting cage bedding. Neutrophil-depleted mice were infected with 10^8^ *C. auris* by tail vein injection. (A) Before autoclaving, swabbed bedding is shown to have *C. auris* growth on Sabouraud dextrose agar plates, confirmed by PCR. (B) Bedding autoclaved in micro-isolator cages as a closed unit displays no *C. auris* growth, confirmed by PCR.

**Table 2.**
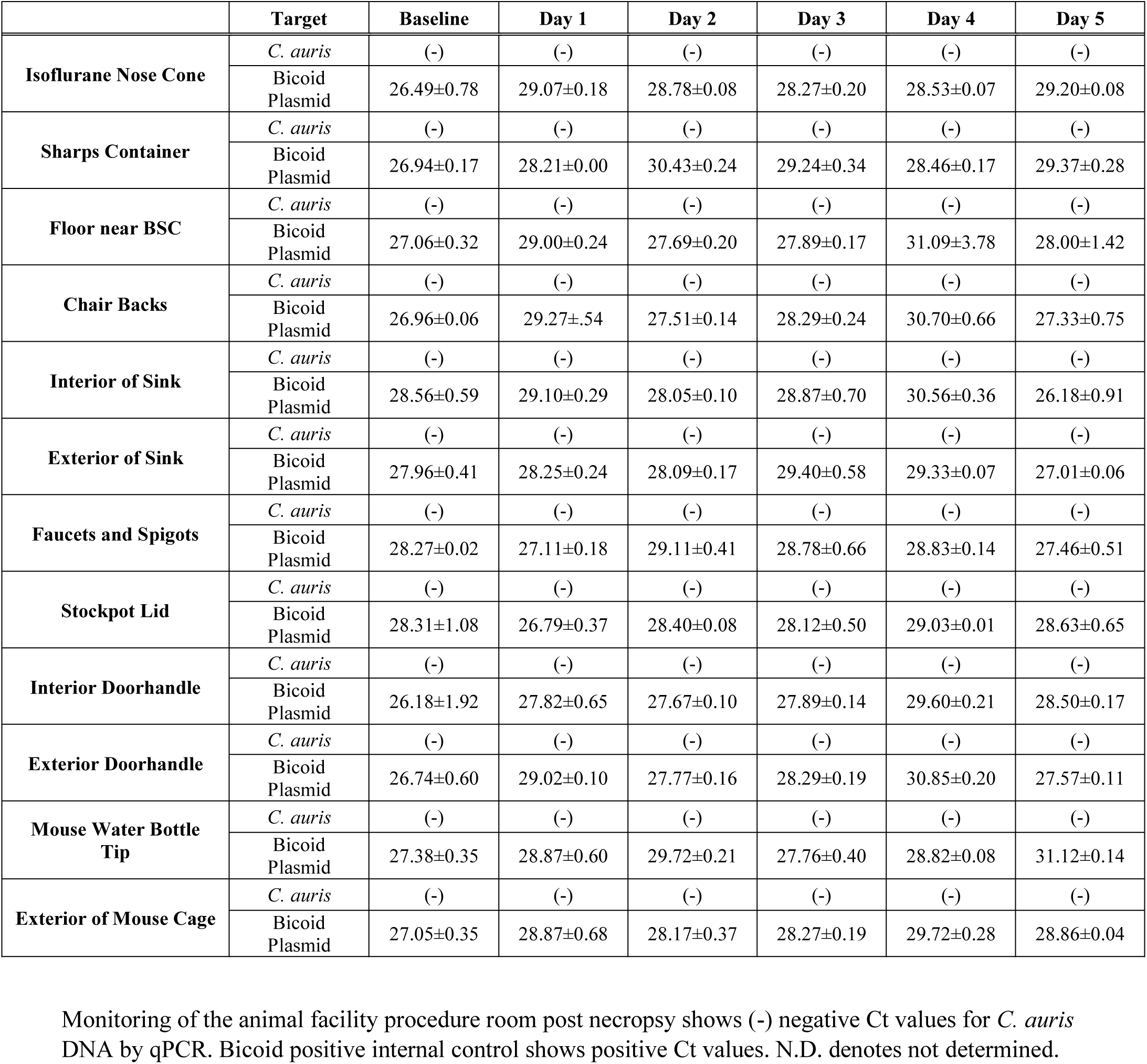
Monitoring of animal procedure room by qPCR.

**Table 3.** Monitoring inside the biosafety cabinet at different inoculums by qPCR.

|  | Target | 1x10<sup>5</sup> | 1x10<sup>6</sup> | 1x10<sup>7</sup> | 1x10<sup>8</sup> |
| --- | --- | --- | --- | --- | --- |
| **Mouse Bedding**<br>**(Before Autoclave)** | *C. auris* | (-) | (-) | (+)<br>29.52±0.16 | (+)<br>28.90±0.04 |
|  | Bicoid Plasmid | 28.38±0.11 | 27.66±0.53 | 26.80±0.28 | 27.42±0.98 |
| **Mouse Bedding**<br>**(After Autoclave)** | *C. auris* | ND | ND | ND | (-) |
|  | Bicoid Plasmid | ND | ND | ND | 24.93±0.33 |
| **Exterior Gloves**<br>**(Before Disinfection)** | *C. auris* | (-) | (+)<br>27.65±0.10 | (+)<br>28.36±0.19 | (+)<br>29.01±0.12 |
|  | Bicoid Plasmid | 28.21±0.23 | 28.34±0.07 | 27.22±0.28 | 27.58±0.27 |
| **Exterior Gloves**<br>**(After Disinfection)** | *C. auris* | (-) | (-) | (-) | (-) |
|  | Bicoid Plasmid | 27.65±0.26 | 28.74±0.42 | 29.72±0.11 | 28.96±0.66 |
| **Interior of BSC**<br>**(Before Disinfection)** | *C. auris* | (-) | (-) | (-) | (+)<br>22.93±0.20 |
|  | Bicoid Plasmid | 26.90±0.02 | 28.20±0.01 | 28.64±0.24 | 34.10±4.65 |
| **Interior of BSC**<br>**(After Disinfection)** | *C. auris* | (-) | (-) | (-) | (-) |
|  | Bicoid Plasmid | 28.90±0.51 | 27.93±0.23 | 28.34±0.02 | 27.36±0.47 |

Monitoring within the biosafety cabinet within the procedure room post necropsy across several concentrations shows (+) positive Ct values for *C. auris* DNA by qPCR. Monitoring within the biosafety cabinet post-disinfection and post-autoclaving samples shows (-) negative detection of *C. auris* DNA by qPCR. Bicoid positive internal control shows positive Ct values. N.D. denotes not determined.

Positive tests of the bedding at 10^7^ and 10^8^ inoculums provided the opportunity to test whether autoclaving and disinfection were efficient. To remediate *C. auris* in the bedding, the closed micro-isolator cages with water bottle attached were autoclaved at 121°C for 60 min. Both culture and PCR results were negative, revealing that autoclaving is essential for cage disinfection. At 10^7^ and 10^8^ inoculums, oral and anal swabs were taken from mice to determine if these were the source of *C auris*. Swabs were cultured in three different media, including (1) Sabouraud Dextrose Agar plates with antibiotics, (2) Sabouraud Dulcitol Agar plates with 10% NaCl and antibiotics, and (3) Sabouraud Dulcitol, 10% NaCl and antibiotics broth. The cultures did not show any growth at these concentrations after two weeks; therefore, no PCR was performed on these swabs (data not shown).

During necropsies, the interior of the biosafety cabinet was only positive for *C. auris* inoculums of 10^8^ cells. The disinfection protocol of 10% bleach for five min followed by 70% ethanol was found to be effective, as the interior of the biosafety cabinet tested negative for *C. auris* by PCR and by culture following this procedure. Glove exteriors during necropsies were positive at 10^6^, 10^7^ and 10^8^ concentrations, with Ct counts of 27.65, 28.36 and 29.01, respectively, indicating that this an important potential source of contamination in the procedure room. To easily remediate this potential source of contamination, exterior gloves were removed right after the full necropsy of each mouse was done and before the workers’ hands touched anything else within the biosafety cabinet. As an important reminder, the “buddy” always kept a watch for the removal of exterior gloves.

### Post-Six-Month Monitoring of the Procedure Room Using *C. auris* Inoculums Ranging from 10^5^-10^8^ Cells

To assess the effectiveness of the workflow and disinfection protocol, the procedure room was assessed six months after initial studies began. Throughout this time, at least twenty experiments were done at inoculums ranging from 10^5^-10^8^. Areas swabbed included the isoflurane nose cone, sharps container, the floor under the biosafety cabinet, the backs of the chairs, the interior and exterior of the sink, the facets and spigots of the sink, the stockpot lid, and the interior and exterior door handles, as well as the mouse restrainer. We swabbed these areas in replicate to ensure that there were no false negatives. The results summarized on Table 4 showed all negative Ct values for all of the areas sampled, providing a strong indicator that the workflow and disinfection protocol was effective.

**Table 4.** Monitoring of the procedure room six months later.

|  | Target | Ct Value |
| --- | --- | --- |
| Isoflurane Nose Cone | *C. auris* | - |
|  | Bicoid Plasmid | 27.89±0.44 |
| Sharps Container | *C. auris* | - |
|  | Bicoid Plasmid | 27.80±0.22 |
| Floor near BSC | *C. auris* | - |
|  | Bicoid Plasmid | 28.32±0.06 |
| Chair Backs | *C. auris* | - |
|  | Bicoid Plasmid | 28.08±0.46 |
| Interior of Sink | *C. auris* | - |
|  | Bicoid Plasmid | 29.88±3.81 |
| Exterior of Sink | *C. auris* | - |
|  | Bicoid Plasmid | 27.41±0.02 |
| Faucets and Spigots | *C. auris* | - |
|  | Bicoid Plasmid | 28.04±0.33 |
| Stockpot Lid | *C. auris* | - |
|  | Bicoid Plasmid | 27.96±0.44 |
| Interior Doorhandle | *C. auris* | - |
|  | Bicoid Plasmid | 28.64±0.08 |
| Exterior Doorhandle | *C. auris* | - |
|  | Bicoid Plasmid | 28.29±0.21 |
| Mouse Restrainer | *C. auris* | - |
|  | Bicoid Plasmid | 27.83±0.10 |

Monitoring of procedure room six months after working with *C. auris* inoculums that ranged from 10<sup>5</sup>-10<sup>8</sup> cells. All areas show (-) negative Ct values for *C. auris* DNA by qPCR. Bicoid positive internal control shows positive Ct values.

## DISCUSSION

As an MDR emerging pathogen, recent work in the *C. auris* field has focused on several areas such as the development of rapid identification methods to prevent delayed treatment,^(22, 27)^ the epidemiology and genomic analysis of *C. auris,*^(1)^ the testing of disinfectants and best practices to reduce *C. auris* persistence, especially in healthcare settings,^(10)^ and the testing of small molecule antifungal drugs such as APX001 and the echinocandin rezafungin as alternative therapeutics.^(23, 24)^ Because *C. auris* is a new organism, there are a limited number of animal models. The current animal models use immunosuppressive agents such as cyclophosphamide to establish a *C. auris* infection.^(23)^ However, to date no studies have evaluated the risk of *C. auris* as an occupational exposure issue during animal experiments for researchers and animal care staff.^(23)^ As part of this study, rigorous safety procedures were developed, and *C. auris* was handled at Animal Biosafety Level 2 (ABSL-2) with Animal Biosafety Level 3 (ABSL-3) containment practices that included enhanced Personal Protective Equipment (PPE) and a strict no open handling policy. A “buddy system” was also implemented in the workflow to prevent accidental contamination of the room, as well as allowing researchers to maintain clearly defined clean and dirty work areas. The “buddy system” was also a way for workers to monitor each other and ensure that exterior PPE was changed during cage changes, infections and necropsies. Since exterior gloves tested positive at inoculums of 10^6^-10^8^ cells, they were considered to be important sources of contamination, emphasizing the importance of changing exterior gloves even when moving within the biosafety cabinet. Another important part of the developed procedures included keeping mice in micro-isolator cages, as *C. auris* was detected as positive at 10^7^ and 10^8^ inoculums. The closed micro-isolator cages were effective at keeping dirty bedding isolated and contained water bottles that could be autoclaved as a single unit. This system prevented exposure of animal caretakers to *C. auris* positive bedding when handling cages or if cages fell to the floor prior to autoclaving. Autoclaving was also found to be highly effective in the elimination of *C. auris* in the bedding. This study and others also found the use of 10% bleach followed by 70% ethanol to be an effective disinfectant regimen against *C. auris*,^(4)^ as mice were clearly infected, yet all surfaces were qPCR-negative for *C. auris*.

## CONCLUSIONS

Because there are currently a limited number of documented murine models of *C. auris* infection, occupational exposure of working with these models has not been clearly defined. This study tested both the animal holding room and procedure room and evaluated potential sources of contamination and exposure during animal infections. Using an intravenous model of *C. auris* infection, shedding of the organism was determined to be dose-dependent. *C. auris* was detectable in the cage bedding when mice were infected with 10^7^ and 10^8^ cells, but not with doses of 10^5^ and 10^6^ cells. Testing by anal and oral swabs on mice was unable to identify the source of shedding. Results indicated potential for exposure with *C. auris* during necropsies and when working with infected tissues.

To mitigate these potential exposures, laboratory personnel used two layers of PPE and the lab developed a rigorous “buddy system” workflow. The “buddy’s” role was to remind the person working inside of the biosafety cabinet to change their exterior gloves, ensure that there were no needlestick accidents during infection, and enable the person to move around the procedure room and hand over cages and materials to the person working inside of the biosafety cabinet. The rigorous use of 10% bleach and 70% ethanol and the use of closed micro-isolator cages and cage autoclaving were all determined to be essential. Assessment of the procedure room after six months of experiments using this workflow and disinfection protocol found negative Ct values, indicating that these methods were successful. In conclusion, this study showed that it is possible to work with large inoculums of *C. auris* in an animal facility, and that by following these recommendations, it can be done safely without exposing the researchers and animal caretakers to infection.

## RECOMMENDATIONS

For any study using a *C. auris* infection models, it is recommended that investigators handle the organism at Animal Biosafety Level 2 (ABSL-2) with Animal Biosafety Level 3 (ABSL-3) containment practices. In addition, researchers should wear two layers of protective PPE, such as double gloves, booties, and lab coats. Also, the “buddy system” and a directional workflow should be used, and micro-isolator cages should be used and autoclaved. To maintain a clean animal holding and procedure rooms, surfaces should be decontaminated with 10% bleach for five minutes followed by 70% ethanol. Following these recommendations will ensure a safe laboratory environment for both the researchers and animal care takers.

